# Wolverine density distribution reflects past persecution and current management in Scandinavia

**DOI:** 10.1101/2022.11.14.516397

**Authors:** Ehsan Moqanaki, Cyril Milleret, Pierre Dupont, Henrik Brøseth, Richard Bischof

**Affiliations:** Faculty of Environmental Sciences and Natural Resource Management, Norwegian University of Life Sciences, P.O. Box 5003, 1432 Ås, Norway; Department of Terrestrial Ecology, Norwegian Institute for Nature Research, PB 5685 Torgarden, 7485 Trondheim, Norway

**Keywords:** Abundance, Density, Distribution, Large carnivores, Non-invasive monitoring, Spatial capture-recapture, Transboundary wildlife, *Gulo gulo*

## Abstract

After centuries of intense persecution, several large carnivore species in Europe and North America have experienced a rebound. Today’s spatial configuration of large carnivore populations has likely arisen from the interplay between their ecological traits and current environmental conditions, but also from their history of persecution and protection. Yet, due to the challenge of studying population-level phenomena, we are rarely able to disentangle and quantify the influence of past and present factors driving the spatial distribution and density of these controversial species. Using spatial capture-recapture models and a data set of 742 genetically identified wolverines *Gulo gulo* collected over ½ million km^2^ across their entire range in Norway and Sweden, we identify landscape-level factors explaining the current population density of wolverines in the Scandinavian Peninsula. Distance from the relic range along the Swedish-Norwegian border, where the wolverine population survived a long history of persecution, remains a key determinant of wolverine density today. However, regional differences in management and environmental conditions also played an important role in shaping spatial patterns in present-day wolverine density. Specifically, we found evidence of slower recolonization in areas that had set lower wolverine population goals in terms of the desired number of annual reproductions. Management of transboundary large carnivore populations at biologically relevant scales may be inhibited by administrative fragmentation. Yet, as our study shows, population-level monitoring is an achievable prerequisite for a comprehensive understanding of the distribution and density of large carnivores across an increasingly anthropogenic landscape.

## 1 Introduction

Species distributions we observe today are the result of not only ecological traits and current local environmental conditions, but also land-use history, human activity, and management strategies (Donohue et al. 2000, Foster et al. 2003, Di Marco and Santini 2015). Emerging disturbance regimes, such as altered frequency and intensity of extreme weather and climate events (Ummenhofer and Meehl 2017), further impact species distributions. Identifying and disentangling the factors that lead to the distribution and dynamics of species is one of the most profound and long-standing research areas in ecology, with both fundamental and applied implications (Guisan and Zimmermann 2000, Elith and Leathwick 2009, Jetz et al. 2019).

Humans are the main transformers of Earth’s ecosystems (Ellis 2011, Pereira et al. 2012, Waters et al. 2016), with a growing list of documented effects on wildlife (Yackulic et al. 2011, Tucker et al. 2018). Despite a broad overall consistency in wildlife responses to anthropogenic disturbances, there is considerable variability in scale, magnitude, and pattern of human impacts (Tablado and Jenni 2017, Gaynor et al. 2018, Tucker et al. 2018). A popular example is the case of large carnivore species that have undergone substantial range contractions due to intensive persecution by humans. While many species continue to struggle, some have in recent decades successfully recolonized part of their historic range, particularly in Western Europe and North America (Linnell et al. 2001, Zedrosser et al. 2011, Chapron et al. 2014, Ripple et al. 2014, Ingeman et al. 2022). Limited understanding of factors shaping the spatial configuration of carnivore populations poses a challenge to science and management, and the current knowledge gaps may hinder predictions of future responses in the face of increasing human pressure.

The fall and rise of wolverines *Gulo gulo* in Scandinavia is a prime example of recovery of an iconic large carnivore following intense persecution and range contraction. The wolverine was historically distributed throughout most of the Scandinavian Peninsula (Landa et al. 2000, Flagstad et al. 2004). During the twentieth century, intensive persecution of the wolverine reduced its range and population size drastically. By 1970, the population was functionally extinct in many areas with the exception of a narrow strip in the alpine region along the border between Sweden and Norway (Landa et al. 2000, Flagstad et al. 2004; Fig. 1). The situation was similarly grim in neighboring Finland, where wolverine observations were rare beyond the borderland with Russia (Lansink et al. 2020; Fig. 1). The wolverine finally received legal protection in both Norway and Sweden by 1973, and later followed by Finland, and gradually recolonized many parts of its historical range in Scandinavia (Flagstad et al. 2004, Aronsson and Persson 2017, Lansink et al. 2020). Today, the wolverine population is established across Norway and Sweden beyond the alpine refuge areas (Chapron et al. 2014, Gervasi et al. 2016, Bischof et al. 2020). The return of the wolverine has rekindled conflict with the sheep-farming industry and semi-domesticated reindeer *Rangifer tarandus* husbandry (Flagstad et al. 2004, Hobbs et al. 2012, Persson et al. 2015, Aronsson and Persson 2017). The wolverine is listed on Appendix S2 of the Bern Convention for both countries and is therefore formally “strictly protected”. However, because Norway is not a member of the European Union, it is not bound by the same set of regulations. Wolverines are therefore subject to persistent lethal control in Norway, while they are strictly protected in Sweden under the European Union’s Habitats Directive 92/43 (annex IV; Habitats Directive 1992).

**Figure 1:**
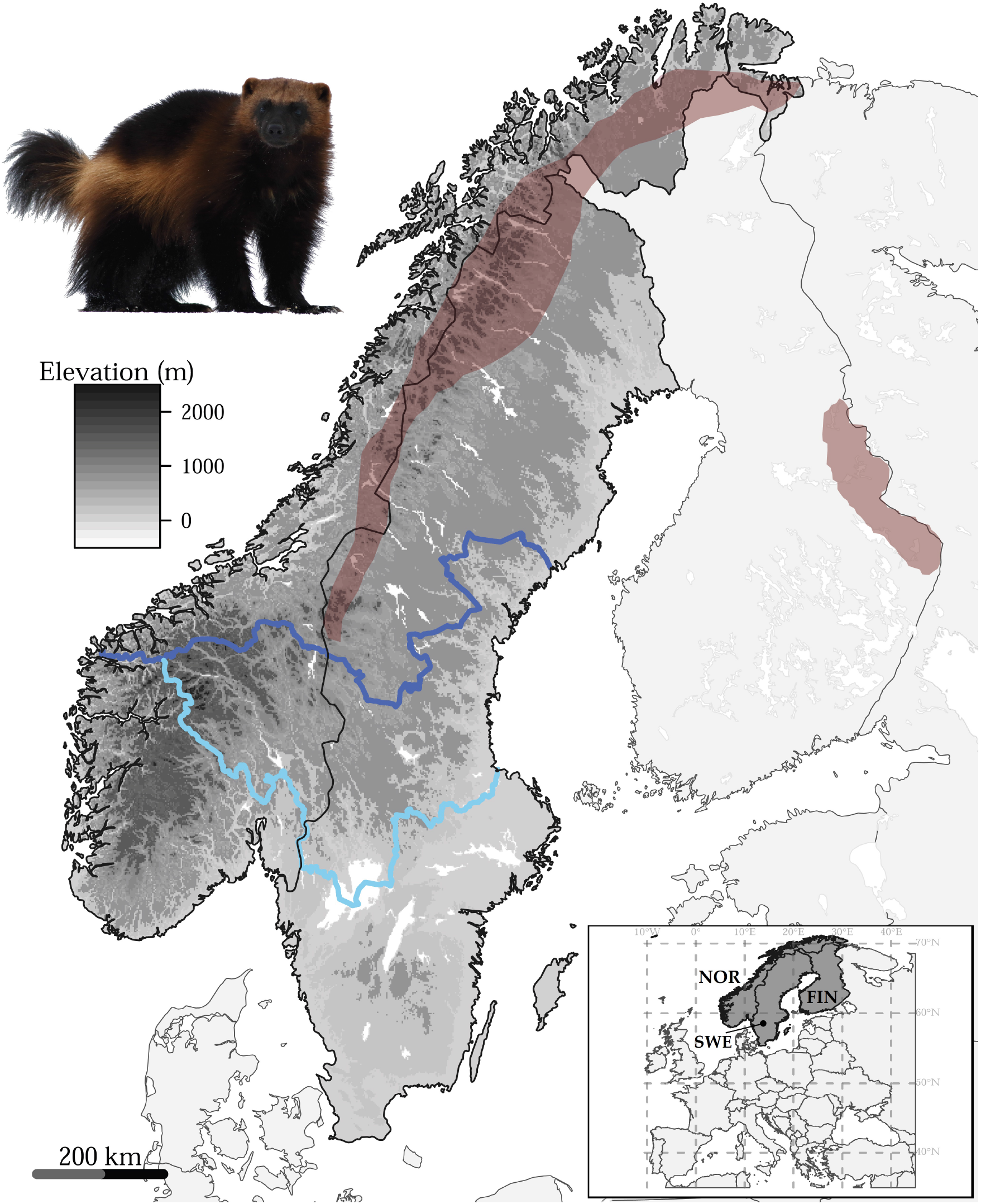
Approximate wolverine *Gulo gulo* distribution in Scandinavia (red polygon on the left) and Finland (red polygon on the right) in the 1970’s, when the population range was at its lowest in modern times following intense human persecution (i.e., the relic range; redrawn after Landa et al. (2000) and Chapron et al. (2014)). Blue lines separate zones containing administrative units (i.e., large carnivore management regions in Norway and counties in Sweden) with shared population goals for the wolverine (see Table 1). We merged the zones below the dark blue line into one southern zone in each country. Photo credit: Karel Bartik/www.shutterstock.com

In a human-dominated world, understanding population-level drivers of species spatial distribution and particularly density is important to understand and predict the potential for species-environment interactions in a management context. What we know about landscape and environmental factors influencing wolverine distribution and density has been cobbled together from a small patchwork of studies, often with limited spatial extent, in various parts of the global distribution range of the species (Fisher et al. 2022). In Scandinavia, population and landscape-level determinants of wolverine distribution and density are poorly known. Historical (Landa et al. 2000) and current (Chapron et al. 2014) range maps suggest that recolonization in this anthropogenic landscape has been facilitated by favorable legislation and improved cultural acceptance (Linnell et al. 2001, Flagstad et al. 2004, Aronsson and Persson 2017). However, there is evidence that biophysical constraints, such as climate, habitat, and terrain, have played a greater role in shaping the current spatial distribution of the wolverine at the continental scale (Cretois et al. 2021). Current management decisions use information that is largely based on data from the high-conflict alpine areas (Brøseth et al. 2010, Aronsson and Persson 2017) but would benefit from a better knowledge of the determinants of wolverine’s spatial variation in density across its entire Scandinavian range. Until recently, this was out of reach because of the rarity and elusive behavior of the species, the vast geographic expanse of the population, and spatially incomplete surveys (Flagstad et al. 2004, Gervasi et al. 2016, Aronsson and Persson 2017).

Here, we set out to quantify the extent to which current wolverine population density across the Scandinavian Peninsula is affected by past and present conditions. Importantly, we do so for the entire ½ million km^2^ range of the species across Norway and Sweden. Three major challenges plague monitoring of elusive species, such as the wolverine, at ecologically relevant scales: (1) the collection of sufficiently detailed individual data from an entire population, (2) imperfect detection (i.e., not all individuals in the population are detected), and (3) a paucity of computationally efficient analytical tools to disentangle the effects of ecological drivers from both stochastic process noise and observation errors (Isaac et al. 2020, Cretois et al. 2021, van de Schoot et al. 2021). In this study, we tackled these challenges for the Scandinavian wolverine by analyzing a comprehensive capture-recapture data set of genetically identified wolverine individuals across the entire population in Norway and Sweden using recently developed efficient spatial capture-recapture (SCR) models (Bischof et al. 2020, Turek et al. 2021).

## 2 Methods

### 2.1 Non-invasive genetic sampling

We used wolverine non-invasive genetic sampling (NGS) data from the Scandinavian large carnivore monitoring database (Rovbase 3.0; www.rovbase.no and www.rovbase.se). This is one of the largest, long-term capture-recapture data of terrestrial wildlife globally (Bischof et al. 2020, Tourani 2022). Wildlife authorities and volunteers conduct both structured searches and opportunistic sampling of putative wolverine scats and hair on snow between December and June each year throughout the species’ range in Norway and Sweden. The structured search tracks and locations of non-invasive samples are GPS recorded (Fig. S1). Further details on wolverine NGS is provided elsewhere (e.g., Brøseth et al. 2010, Gervasi et al. 2016, Bischof et al. 2020). Samples were processed and analyzed by two dedicated DNA labs using a number of control measures to minimize genotyping errors, as described elsewhere (Ekblom et al. 2018, Flagstad et al. 2019, Lansink et al. 2022). First, samples were analyzed with a Single Nucleotide Polymorphism (SNP)-chip with 96 markers and, second, all individuals were analyzed with 19 microsatellite markers to determine species and identity of wolverine individuals as well as their sex. We used NGS data collected between 1 December 2018 and 30 June 2019, which consisted of individual identity, sex, collection date, and coordinates associated with each wolverine sample. This sampling period represents the latest, most complete, semi-systematic wolverine NGS effort across the entire range of the wolverine population in Scandinavia to date (Flagstad et al. 2019, Bischof et al. 2020, Milleret et al. 2022).

### 2.2 Analysis

SCR models offer a flexible framework to account for imperfect detection of individuals and provide spatially explicit estimates of abundance (i.e., density) and other population parameters (Efford 2004, Borchers and Efford 2008, Royle et al. 2014). The SCR modeling framework can support flexible sampling configurations and incorporate both individual- and detector-level covariates to account for sources of heterogeneity in detectability, and spatial covariates to account for variation in density (Royle et al. 2014). Although building spatially indexed hierarchical models, such as SCR, can be computationally challenging or even prohibitive for large spatial extents, recent developments have resulted in dramatic improvements (e.g., Milleret et al. 2019, Turek et al. 2021, Zhang et al. 2022). Here, we build on these recent developments to study the landscape-scale determinants of the Scandinavian wolverine density.

#### 2.2.1 Spatial capture-recapture model

We built a single-season (i.e., demographically closed) SCR model in a Bayesian framework by expanding on our previous work (Bischof et al. 2020). Our SCR model contains two hierarchical levels: (1) The observation sub-model accounts for imperfect and variable wolverine detectability during NGS; and (2) The ecological sub-model describes wolverine density as the main ecological process of interest in this study. Our SCR model estimates the following parameters: (1) the baseline detection probability *p*_0_: detection probability at a trap or hypothetical detector located at an animal’s activity center *s_i_,* a latent variable representing the expected location about which an individual uses space during the sampling period; (2) the spatial scale parameter of the detection function *σ;* (3) the number *N* of wolverine activity centers within the available habitat *S* (i.e., the detector grid and a buffer around it; see below), which can be used to derive density *D* (see below); and (4) the effects (regression coefficient *β*) of spatial and individual covariates on the detection probability and density.

##### (1) The observation sub-model

We used the conventional half-normal detection function (Borchers and Efford 2008, Royle et al. 2014) to model the probability *p* of detecting individual *i* at detector *j* as a decreasing function of the distance *d* between the detector and the individual’s center of activity 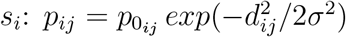. The detection function is assumed to reflect individual space use and is therefore directly linked with the home range concept (Royle et al. 2014). Because we used a data-augmentation approach (Royle et al. 2007), the detection of an individual has to be made conditional on the individual’s state *z_i_* (*z_i_* = 1 when individual *i* is member of the population *N*), which is governed by the inclusion probability *ψ*: *z_i_* ~ Bernoulli(*ψ*). The population size can be then derived by summing the *z_i_*’s: 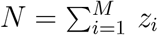, where *M* is the chosen size of the data-augmented population (Royle et al. 2007) and represents the maximum number of wolverines in the habitat *S* (see *Ecological sub-model*).

In our study, detectors are the centers of 5 572 10 × 10 km grid cells, covering a land area extending 100 km beyond the outermost wolverine NGS detections collected during the sampling period (Fig. S1). We used a partially aggregated binomial observation model (Milleret et al. 2018) to retain more information from the wolverine NGS data by dividing each main detector cell into 25 sub-detector cells of 2 × 2 km. By retrieving the number of sub-detector cells with at least one non-invasive sample for each wolverine detected at each main detector cell, we generated individual spatial detection histories (Royle et al. 2014). Finally, we placed a 40-km buffer around the detector grid to define the habitat *S*. This value was chosen based on the average home-range radius of adult Scandinavian wolverines (Persson et al. 2010, Mattisson et al. 2011, Aronsson et al. 2022), so that the buffer is larger than three times the estimated *σ* of 10.3 km (95% credible interval [CI] = 10.1 – 10.5 km) for male wolverines as reported by Bischof et al. (2020). This buffer area allows detection of individuals even if their activity centers are located outside the detector grid (Efford 2004, 2011). The detector grid covered most of the contiguous Scandinavian Peninsula over Norway and Sweden (58° 08’ - 70° 42’ N, 5° 56’ - 32° 46’ E; Fig. S1), while parts of the buffer (41.6%) fell inside Finland and Russia. Thus, the available habitat was 633 200 km^2^, after removing large lakes and other non-contiguous land areas, of which 88% (557 200 km^2^) were in Norway and Sweden (Fig. S1).

Wolverine NGS was conducted by hundreds of field staff and volunteers across different jurisdictions in Norway and Sweden. We therefore expected spatial variability in detection probability of wolverine individuals (Efford et al. 2013, Moqanaki et al. 2021). Following Bischof et al. (2020), we considered a different baseline detection probability for each jurisdiction *p*_0_*County*__ (*County* = 1:8) to account for possible regional differences in monitoring regimes. Jurisdictions were defined based on carnivore management regions in Norway and counties in Sweden after Bischof et al. (2020) with slight modifications to match with our habitat extent (Fig. S2). We merged neighboring jurisdictions to ensure sufficient wolverine detections for estimating baseline detection probability in each unit (Bischof et al. 2020). In addition, we modeled the effect of three detector- and one individual-level covariates that may influence the probability of wolverine detection (Table S1):

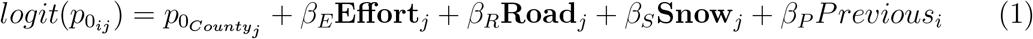

**Effort**_*j*_ is the length (m) of GPS search tracks within each detector grid cell *j* recorded during the structured NGS, **Road**_*j*_ is the logarithm of the average geographic distance (km) from each detector to the nearest road of any type, and **Snow**_*j*_ is the average percentage of snow cover in each detector grid cell during the sampling months (December 2018 - June 2019; Table S1). We also modeled individual variation linked with detection in the previous sampling year *Previous_i_*; a binary covariate which takes the value 1 if individual *i* was detected in the previous sampling year and 0 otherwise. During NGS, investigators are believed to have the tendency to prioritize searching in locations where their searches were previously successful, which could positively influence the detection probability of those previously-detected wolverine individuals during the focal sampling year (Gervasi et al. 2014, Milleret et al. 2022). Availability of the monitoring data from the previous year made it possible to account for this potential source of heterogeneity in wolverine detectability. This individual binary covariate *Previous_i_* is latent for augmented individuals and was modeled following a Bernoulli distribution: *Previous_i_* ~ *Bernoulli*(*π*), where *π* is the probability that an arbitrary individual from the population was detected in the previous year. All continuous spatial covariates were scaled before SCR model fitting. Further details on detection covariates, the rationale to include them, and their original source and spatial depiction are provided in Table S1 and Figure S3.

##### (2) The Ecological sub-model

describes the number and distribution of all wolverines present in the population (i.e., detected and non-detected). We used a data augmentation approach (Royle et al. 2007) to account for those wolverine individuals that were not detected during NGS, where the super-population size *M* (i.e., detected and augmented individuals) is chosen to be considerably larger than *N*. Following Bischof et al. (2020) and given the relatively high detectability of the target population during NGS (Milleret et al. 2022), we chose an augmentation factor of 0.8 to facilitate the analysis by Markov chain Monte Carlo (MCMC). Thus, *M* was large enough such that the probability that *M* individuals were alive in *S* during NGS was negligible.

SCR estimates of abundance are spatially explicit, meaning that they are derived from the estimated location of all individual activity centers *s_i_* with *z_i_* = 1 across the available habitat *S* (Efford 2004, Borchers and Efford 2008, Royle et al. 2014). The collection of activity centers can be seen as the realization of a statistical point process (Illian et al. 2008). To study how wolverine density varies in Scandinavia in response to a number of environmental and history-related covariates (Table 1), we used an inhomogeneous binomial point process to model spatial variation in the distribution of individual activity centers with intensity function (Zhang et al. 2022): *λ*(*s*) = *e*^*β***X**(*s*)^, where **X**(*s*) is a vector of spatial covariate values evaluated at location *s* and *β* is a vector of associated regression coefficients. The intensity function *λ* conditions the placement of activity centers within each of the 20 × 20 km habitat grid cells *s* used in this analysis (Fig. S1). In this formulation, no intercept is needed as the number of activity centers is conditioned by data augmentation; thus, regression coefficients represent the relative effects of the different covariates on wolverine density (Zhang et al. 2022).

**Table 1:**
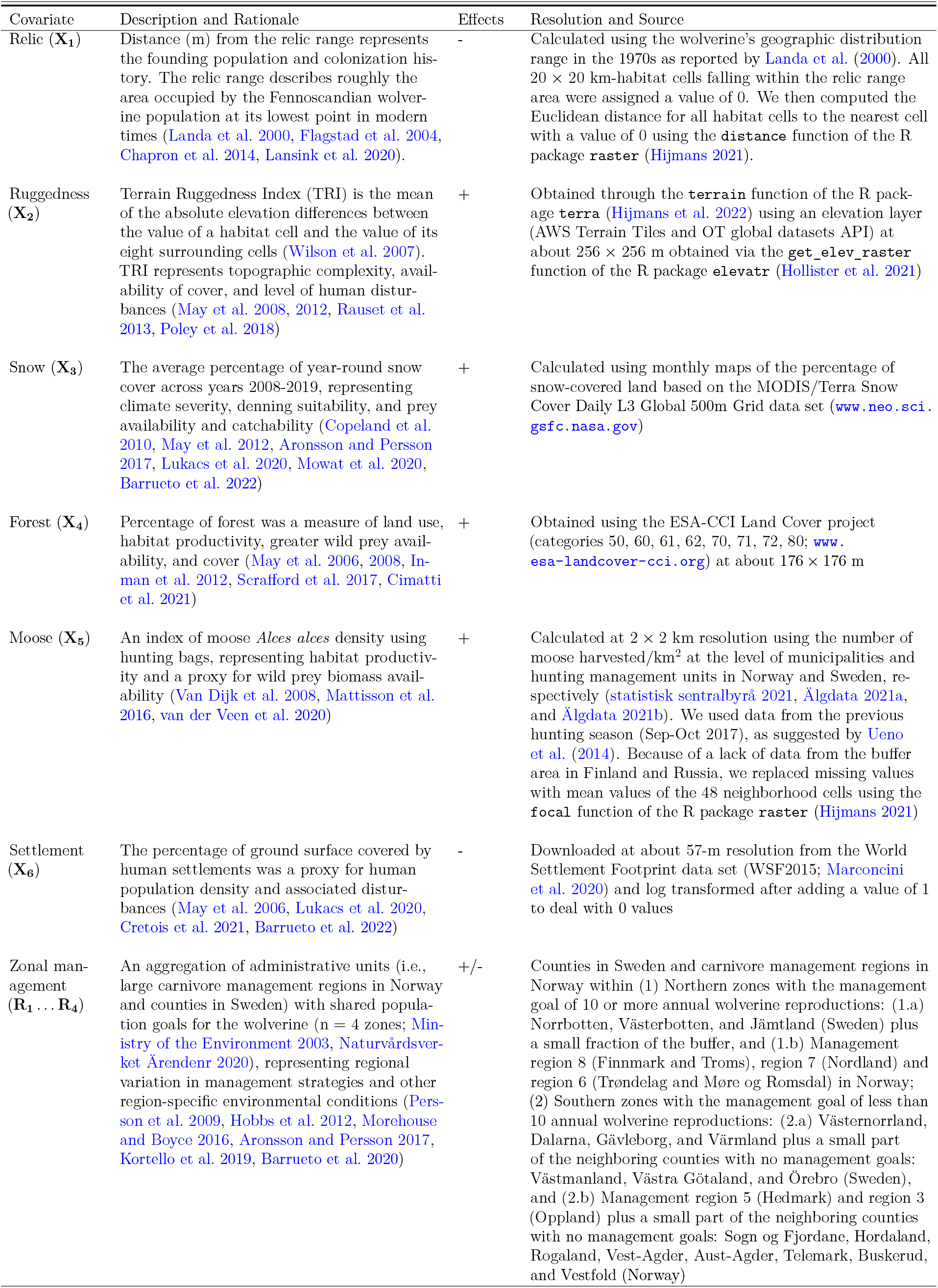
Description, rationale for inclusion, expected effects, and source and native spatial resolution of covariates of density used to model the density distribution of the wolverine *Gulo gulo* across Norway and Sweden between December 2018 and June 2019

To disentangle the determinants of wolverine density within Scandinavia, we measured habitat characteristics at the scale of the home range of a wolverine (i.e., the second order of habitat selection; Johnson 1980). We selected biotic and abiotic covariates following previous studies on wolverine distribution and habitat use and preferences (Fisher et al. 2022 and references therein; Table 1). Specifically, we selected covariates that may explain spatial variation in wolverine density in Scandinavia at broad scale (Table 1; Fig. S4): (1) Distance from the relic range (Landa et al. 2000, Flagstad et al. 2004; Fig. 1) to describe recolonization history; (2) Terrain Ruggedness Index (TRI) explaining general topographic complexity; (3) Average percentage of year-round snow cover as a measure of climate suitability (which was different from the snow covariate used as a detector-level covariate; Table S1); (4) Percentage of forest representing land use and habitat productivity; (5) Moose *Alces alces* harvest density as a proxy for wild prey biomass availability, (6) Percentage of human settlement areas as a measure of human disturbances, and (7) Zonal management to account for regional differences in wolverine management plans and other environmental conditions. The impact of current management was specifically included because of unique management goals for wolverines in different areas of Norway and Sweden (Ministry of the Environment 2003, Naturvårdsverket Ärendenr 2020). Briefly, we divided our habitat layer into northern and southern zones in each country (n = 4; Fig. S4, Table 1) by aggregating jurisdictions with similar management goals for the number of wolverine annual reproduction and other environmental conditions (e.g., climate, prey availability and abundance, and human influence). We simplified the spatial variation in wolverine management by merging several counties or carnivore management regions, and partially included jurisdictions in the southern part of each country without management goals (Table 1; Fig. 1), since these southern counties contained no NGS and wolverine detections in our data set (Fig. S1). Likewise, we merged the buffer area in neighboring Finland and Russia with the northern zones (Fig. S4). We then calculated the proportion overlap between each habitat cell and the resulting four zones to define four spatial covariates (Fig. S4). Because the four proportions sum to one, we did not use the first zone covariate to avoid identifiability issues (i.e., the northern zone in Sweden, zone 1.a in Table 1, was an implicit intercept). Since management goals and other zone-specific characteristics of the biotic and abiotic environment may also have affected the wolverine’s ability to recolonize away from the relic range, we included an interaction term between the distance from the relic range and each of the four zones:

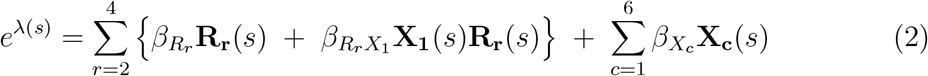

The spatial covariates **X** are the distance from the relic range **X**_1_, Terrain Ruggedness Index **X**_2_, the average percentage of year-round snow cover **X**_3_, the percentage of forest **X**_4_, the percentage of human settlement areas **X**_5_, and the moose harvest density **X**_6_. **R_2_**, **R_3_**, and **R_4_** are the three zone covariates representing southern Sweden and northern and southern Norway (Table 1). In total, we estimated 12 regression coefficients *β*.

We transformed all covariate raster layers from the original projection to the Universal Transverse Mercator (UTM zone 33N) and locally interpolated the raster values using the “bilinear” method of the resample function of the R package raster (Hijmans 2021) to match the 20 × 20 km habitat grid used in this analysis (Figs. S1 and S4). All continuous covariates were then standardized prior to their inclusion in the model to have a mean of zero and one unit standard deviation. Further details regarding the rationale for including each covariate, their sources, and their expected effects are provided in Table 1 and Fig. S4.

#### 2.2.2 Implementation

We fitted SCR models with `NIMBLE` (version 0.12.2; de Valpine et al. 2022) in R (version 4.2.1; R Core Team 2022) for female and male wolverines separately using the recent developments by Turek et al. (2021) and custom functions made available through the R package `nimbleSCR` (Bischof et al. 2021). We ran four MCMC chains, each with 200 000 iterations, discarded the initial 10 000 samples as burn-in, and thinned by a factor of 10. We assessed mixing of chains by inspecting traceplots, and we considered models as converged when the potential scale reduction value 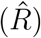 was ≤ 1.10 for all parameters (Brooks and Gelman 1998). Data and R code for fitting the SCR model are provided in the Supplementary Information, and Table S2 shows the list of priors used.

To explore the relative importance of each covariate on density, we incorporated a Bayesian variable selection approach in `NIMBLE` using reversible jump MCMC with indicator variables (Green 1995, O’Hara and Sillanpää 2009). We incorporated an indicator variable *w* associated with each regression coefficient *β* (n = 12; Table S2). Thus, we modified equation (2) to include (*w* = 1) or exclude (*w* = 0) the effect of each coefficient in the presence of other covariate effects in a given posterior draw: *Λ*(*s*) = *e*^*β*_1_*w*_1_**X**_1_(*s*)^+ ⋯ + *β*_p_w_p_**X**_*p*_(*s*). We constrained inclusion of the interaction coefficients to when the corresponding main effects were also included. For inference on the different coefficients, we discarded MCMC draws where *w* = 0.

We calculated the median and the 95% CI limits of the posterior distribution for all parameters, except for abundance, where we reported mean and 95% CI. To obtain total wolverine abundance, we combined *N* estimates of male and female wolverines by merging posterior MCMC samples from the sex-specific SCR models. In both total and sex-specific models, we summed the total number of predicted activity center locations of alive individuals (*z_i_* = 1) within each habitat cell for each iteration of the MCMC chains; thus, we generated a cell-based posterior distribution of abundance that can be viewed also as density. Using this approach, we extracted abundance and density estimates and the associated uncertainty for different spatial units relevant for wolverine management at the country level, besides the total estimates for the entire population in Scandinavia.

We constructed two types of sex-specific density maps: (1) a realized density map based on the posterior location of activity centers as described above, and (2) an expected density map based on the estimated intensity of the density point process per habitat cell of 20 × 20 km and the estimate of population size: 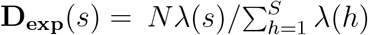. “Realized” density maps show density based on the average model-estimated activity center locations of individuals, as opposed to “expected” density maps, which show predicted density based on the regression model underlying the intensity surface. To present uncertainty, we calculated and mapped the standard deviation of the per-cell posterior of density (Miller et al. 2013).

## 3 Results

### 3.1 Non-invasive genetic sampling

During the sampling period between 1 December 2018 and 30 June 2019, 283 282 km of GPS search tracks were recorded within our designated detector grid (Fig. S1) across Norway (34%) and Sweden (66%). The final NGS data set consisted of 2 444 (1 350 male and 1 094 female) detections from 742 (335 males and 407 females) genetically identified wolverine individuals across the entire population on the Scandinavian Peninsula (Fig. S1). The number of detections (i.e., recaptures) per identified individual ranged from 1 to 13 for both sexes (mean = 3.0 males and 2.1 females).

### 3.2 Density predictors

The variation in wolverine density across Scandinavia was explained by distance from the relic range in different zones, percentage of human settlements, moose harvest density, year-round snow, terrain ruggedness, and percentage of forest (Fig. 2). The magnitude of the effects and uncertainty around them varied moderately between the sexes (Figs. S5 and 2). For both females and males, the effects of being in southern Norway, distance from the relic range in northern Sweden, and percentage of human settlements received the most support based on the inclusion probability (≥ 0.99; Figs. 2 and S5). In addition, for female wolverines the effects of being in northern Norway and distance from the relic range in southern Norway, and for males the effect of moose harvest density received inclusion probabilities of ≥ 0.99 (Figs. 2 and S5).

**Figure 2:**
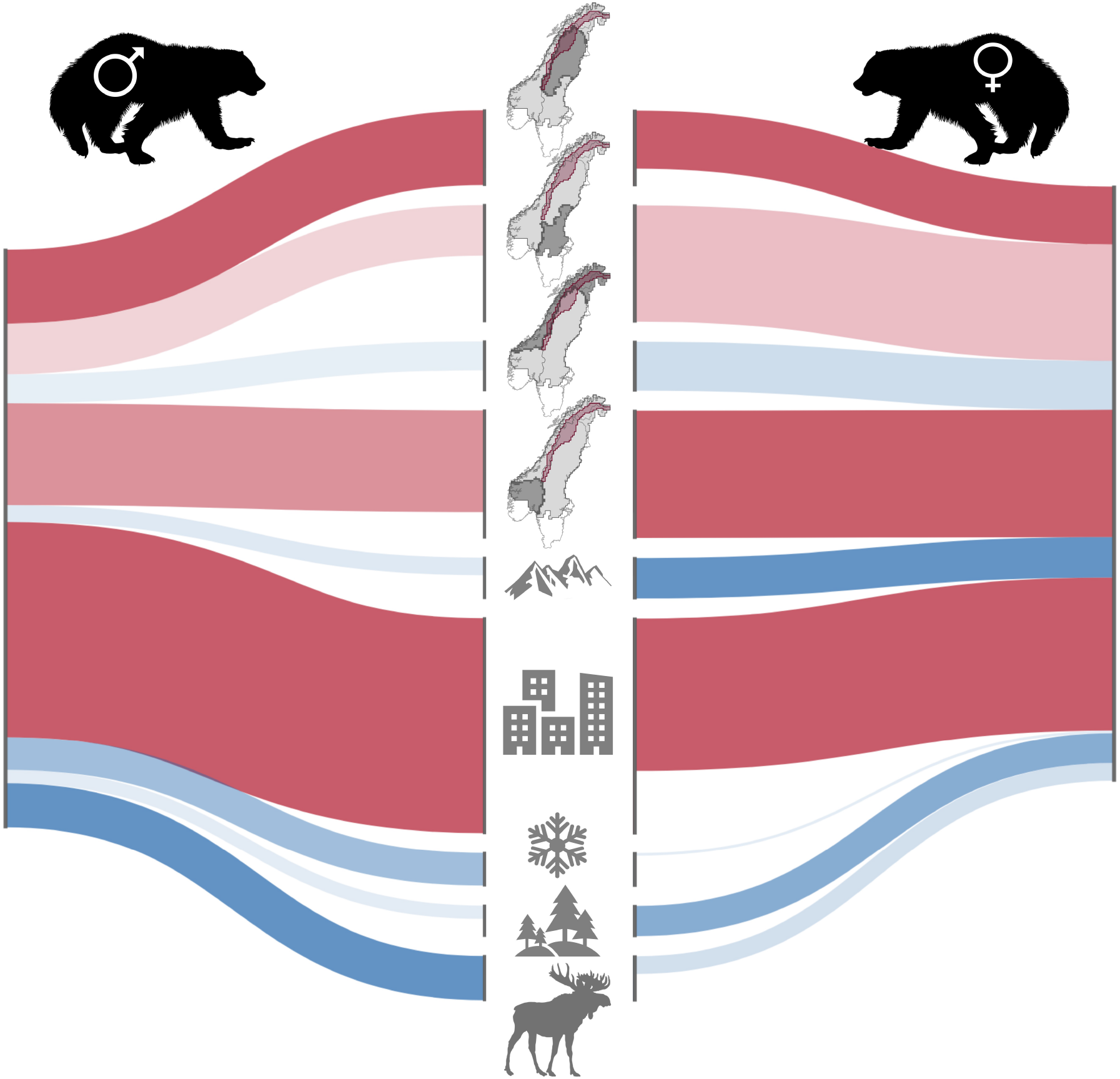
The effect of environmental covariates (middle column) on density of male (left) and female (right) wolverines *Gulo gulo* in Scandinavia between December 2018 and June 2019 as estimated by sex-specific spatial capture-recapture models. Line width represents the magnitude of the median effect (i.e., the thicker, the larger the strength of the covariate effects). Line color shows direction of the effects (blue = positive and red = negative effects), and the opacity level indicates the amount of support for the inclusion of each covariate in the model (inclusion probability of 0 [transparent] to 1 [opaque]). For interaction effects of distance from the relic range in different zones (top four), the line width indicates differences of coefficient estimates from the zone in northern Sweden (the top line). The density covariates are (from top to bottom): Distance from the relic range in (1) northern Sweden, (2) southern Sweden, (3) northern Norway, and (4) southern Norway; (5) Terrain Ruggedness Index; (6) percentage of human settlements; (7) the average percentage of year-round snow cover; (8) percentage of forest; and (9) moose *Alces alces* harvest density (Table 1). The main additive effects of zones are not shown (see Fig. S5).

Among the covariates considered, percentage of human settlements had the largest negative effects on both female and male wolverine densities (median and 95% CI *β*_*X*_5__ = –1.61, –2.66 to –0.79 [female] and –2.27, –3.41 to –1.33 [male]; Figs. S5 and 2). Likewise, distance from the relic range negatively affected the density of both sexes, with significantly stronger effects in southern Norway (*β*_*R*_4_*X*_1__ = –1.35, –1.99 to –0.70 [female] and –1.07, –1.87 to –0.26 [male]) compared to the effect of distance from the relic range in northern Sweden (Figs. S5 and 3). Based on our results, we predicted that areas located 30 km away from the relic range, as-the-crow-flies, would have on average about two-third lower expected wolverine densities in the southern zones of Norway and Sweden compared to the northern zones (Fig. 3). Moose harvest density was positively associated with both female and male wolverine densities (*β*_*X*_6__ = 0.19, 0.02 to 0.35 [female] and 0.46, 0.31 to 0.63 [male]; Figs. S5 and 2). The effects of percentage of forest (*β*_*X*_6__ = 0.32, 0.12 to 0.52) and terrain ruggedness on density was significantly positive for female wolverines only (*β*_*X*_2__ = 0.42, 0.25 to 0.59), while the effect of year-round snow cover was positive for males only (*β*_*X*_3__ = 0.35, 0.11 to 0.56; Fig. S5; Table S3).

**Figure 3:**
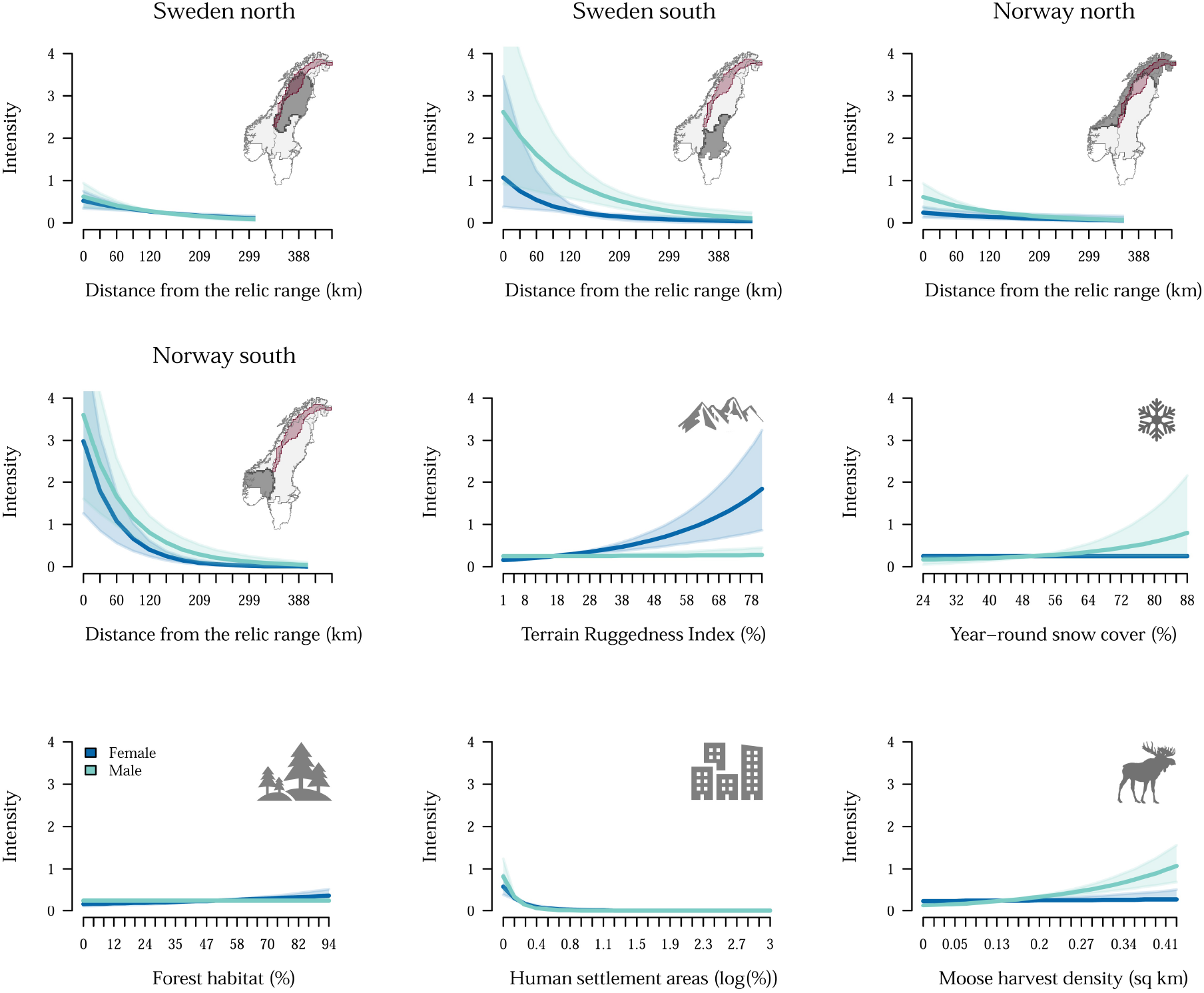
Expected intensity of the density point process for female (blue) and male (green) wolverines *Gulo gulo* in Scandinavia as a function of environmental covariates. Mean response and 95% credible interval are represented by thick lines and transparent polygons, respectively. Predictions in the first four plots from top-left are for the range of values of distance from the relic range (km) that were available in the given zone. The red polygons on the small maps indicate the relic range (Fig. 1) and the dark gray polygons are different zones with contrasting management goals and environmental conditions for the wolverine across the available habitat (Fig. S4).

### 3.3 Detection predictors

The effects of detection covariates varied slightly between male and female wolverines (Table S3). Baseline detection probability *p*_0_ was comparable between sexes (median and 95% CI *p*_0_ = 0.02, 0.01 to 0.02 for both males and females), but varied moderately among the eight carnivore management regions and counties in Norway and Sweden (Fig. S2). Both female and male wolverine detection probabilities increased with search effort (*β_E_* = 0.62, 0.53 to 0.71 [female] and 0.51, 0.44 to 0.59 [male]). Further, for female wolverines, searching farther away from the nearest road increased their detectability (*β_R_* = 0.19, 0.07 to 0.31). Higher percentage of snow cover during the sampling months decreased detectability of males (*β_S_* = −0.22, −0.37 to −0.08). The individual-level covariate representing wolverine detection in the previous sampling year positively influenced male wolverine detectability only (*β_P_* = 0.61, 0.44 to 0.77), suggesting sex-specific detection bias during NGS. The spatial scale parameter was greater for males (*σ_m_* = 8 km, 7.6 - 8.2) than for females (*σ_f_* = 6 km, 5.6 - 6.4). More details are provided in the Supplementary Material.

### 3.4 Sex-specific and total estimates of abundance and density

We estimated the abundance of the Scandinavian wolverine population within our detector grid (Fig. S1) during the 2018/2019 sampling period at 408 (95% CI = 397 - 420) males and 667 (95% CI = 640 - 697) females. The wolverine population in Sweden was estimated to be between 640 and 692 individuals, while in Norway we estimated between 397 and 425 wolverines (Fig S6). Overall, we predicted higher wolverine densities for both males and females closer to the relic range, but the pattern was more pronounced for females (Fig. 4).

**Figure 4:**
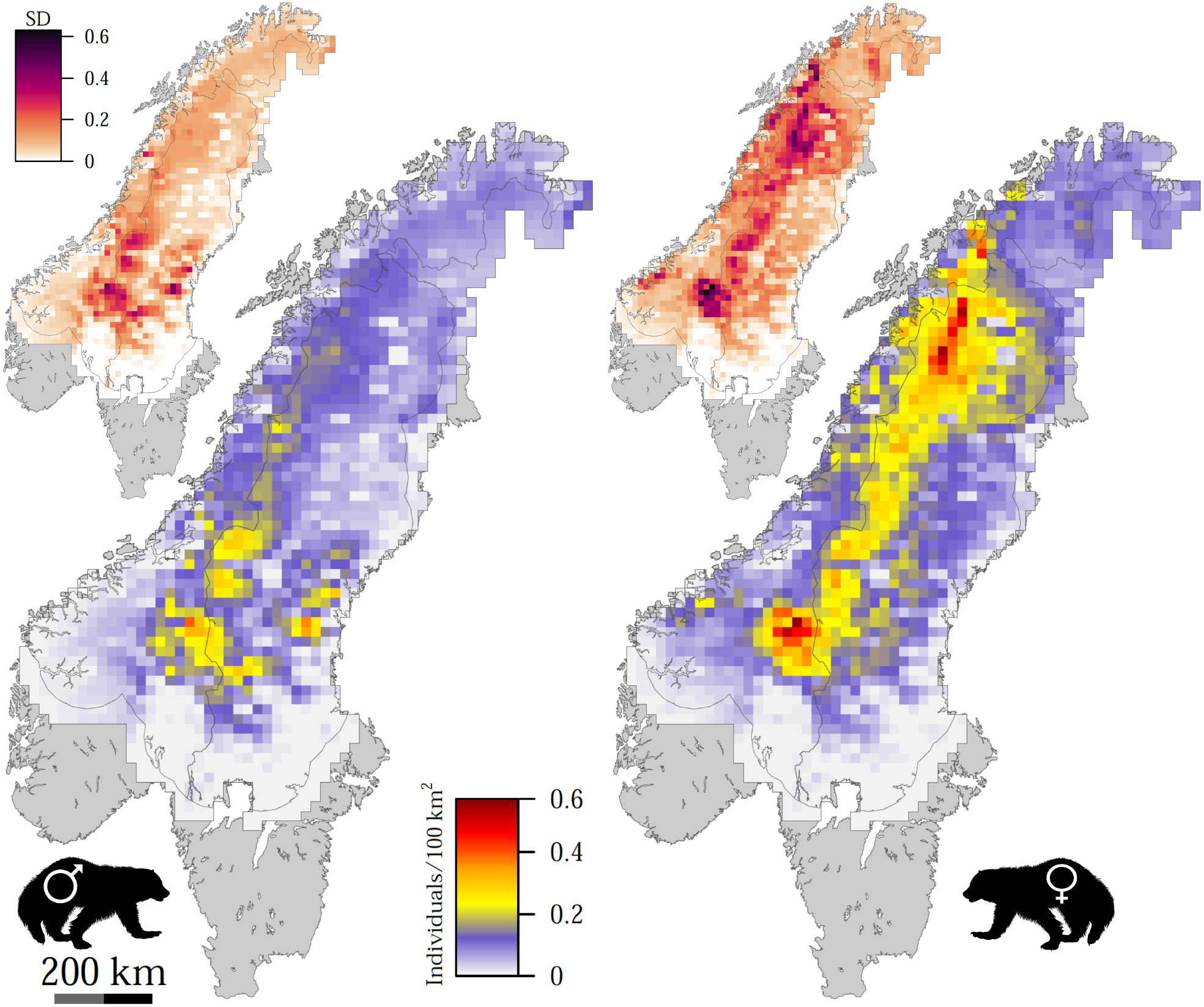
Expected density surfaces of male (left) and female (right) wolverines *Gulo gulo* in Scandinavia as a function of environmental covariates included in spatial capture-recapture analysis (Table 1). The main maps show the average expected density surfaces for each sex (wolverines/100 km^2^) and smaller maps show the standard deviation of predictions.

## 4 Discussion

The present spatial configuration of wolverine density across Scandinavia reflects the species’ recovery from past range-contraction and population decline, modulated by current management goals and environmental conditions. The importance of the relic range along the Swedish-Norwegian border highlights the need for coordinated monitoring and management of this transboundary population of wolverines. Monitoring is already coordinated to some extent (Gervasi et al. 2016, Aronsson and Persson 2017, Bischof et al. 2020), but fully coordinated management is made difficult by existing differences in national and regional population goals and legal obligations.

### The ghosts of the past

A key driver of current wolverine density distribution for both sexes in Scandinavia appears to be distance from the relic range (Figs. 1 and S4), where Scandinavian wolverines survived human persecution before their legal protection in the 1970’s (Landa et al. 2000, Flagstad et al. 2004). We also found that zonal management is one of the main drivers of wolverine density in Scandinavia (Figs. 3 and S5). The density of both male and female wolverines declines with increasing distance from the relic range, and the rate of decline further varies among zones with contrasting management goals regarding wolverine annual reproduction (Figs. 2-3). Regional differences in the effect of distance from the relic range is likely a sign that the current recolonization of wolverines is both a function of past and current management practices and environmental conditions. Together, these factors explained much of the spatial variation in current density of wolverines in the Scandinavian Peninsula (Fig. 4). Whether the relic range represents highly suitable habitat for the Scandinavian wolverine (i.e., historical and current core) or the species was pushed into the alpine refuge areas during the peak of the persecution is not fully understood (Landa et al. 2000, Flagstad et al. 2004, Kerley et al. 2012, Zigouris et al. 2013). Nonetheless, wolverine recolonization in Scandinavia matches the general pattern of return of other large carnivore species in Western Europe and North America (Linnell et al. 2001, Chapron et al. 2014). Successful recovery of these species is partially attributed to changing public attitudes towards large carnivores and effective law enforcement, which, in turn have lowered the risk of direct killing by humans (Zedrosser et al. 2011, Chapron et al. 2014, Ingeman et al. 2022). Likewise, increasing tolerance towards wolverines by Scandinavian farmers and traditional pastoralists has in part been achieved through intensive zonal management of wolverines and compensation schemes (Persson et al. 2015, Aronsson and Persson 2017). Balancing the landscape-level requirements of a viable wolverine (meta-)population and human interests will therefore remain crucial for the successful management.

The ability of wolverines to travel long distances has probably contributed to their successful recolonization in part of their historical range in Scandinavia. However, male wolverines are more likely to disperse, while females usually stay close to their natal range and show high home-range fidelity (Inman et al. 2012, Packila et al. 2017, Aronsson and Persson 2018, Aronsson et al. 2022). We found that spatial covariates tested in our study had qualitatively similar effects on the density of female and male wolverines (Figs. 2 and S5). We note that male and female Scandinavian wolverines have a comparable level of culling mortality (Bischof et al. 2020). Additionally, long-distance dispersal events that lead to successful colonization of unoccupied habitat are not common (Flagstad et al. 2004, Packila et al. 2017). Even if male wolverines on average disperse farther, they may not always successfully establish significantly farther than females. Nonetheless, we observed pockets of higher expected male wolverine density farther from the relic range compared to the expected female density which remained the highest in and near the relic range (Fig. 4). This pattern was reflected in the sex-specific estimates of coefficient for the additive effects of distance from the relic range in the southern zones of Sweden and Norway (Fig. S5).

We estimated, on average, substantially lower wolverine densities in the southern zones of Norway and Sweden compared to the northern zones (Fig. 3). The southern zones generally do not cover semi-domesticated reindeer husbandry areas and calving grounds, but include areas with free-ranging domestic sheep, especially in Norway. The current management strategy in both countries allows more wolverine annual reproduction in the northern zones (Ministry of the Environment 2003, Naturvårdsverket Ärendenr 2020), and the legal removal of wolverines is proportionally more intense in the south to protect the free-ranging sheep (Strand et al. 2019). There are also mismatches between the management goals, their implementation, and regional tolerance of the wolverine in Scandinavia (Aronsson and Persson 2017) that are not entirely reflected by the four zones we considered. Thus, it is likely that the combined effect of the higher cost of dispersal from the relic range and the current management plans regarding wolverine recolonization, together with region-specific environmental characteristics, have resulted in slower wolverine expansion and lower densities in the southern parts of the Scandinavian Peninsula.

### Population-level drivers of variation in density

Wildlife distributions and densities are continuously being shaped by multiple factors at different spatio-temporal scales. Abiotic factors, such as temperature and precipitation, play a key role in shaping species distributions at broad scales (Benton 2009). There is also increasing evidence that biotic factors are important determinants of species distributions at both local and large spatial extents, particularly when accounting for interacting drivers (Van der Putten et al. 2010, Wisz et al. 2013). We found that current environmental features that describe landscape heterogeneity and productivity can explain variation in the Scandinavian wolverine density at the landscape level. Although the relative importance of some of these covariates varied between sexes (Figs. 2 and S5), anthropogenic factors had a consistently negative impact on both male and female wolverine density. Besides quantifying the driving factors of density for the entire population of the Scandinavian wolverines, our study advances the previous findings (Fisher et al. 2022 and references in Table 1) by highlighting the role of past persecution history and current management practices in modulating natural recolonization across a human-dominated landscape.

Human-caused mortality and anthropogenic fragmentation of habitat are limiting wolverine distribution and density globally (May et al. 2006, Persson et al. 2009, Mowat et al. 2020, Lukacs et al. 2020, Lansink et al. 2022, Barrueto et al. 2022). Within the Scandinavian large carnivore guild, wolverines are believed to be the most sensitive to habitat fragmentation (May et al. 2008). We included the percentage of human settlement areas as a measure of human pressure on the natural environment (Marconcini et al. 2020), which represents human population density and the associated disturbances. The negative impact of human settlements on wolverine density appeared to be substantial (Figs. 2 and S5), and we observed drastic declines in the expected density of both male and female wolverines with increasing human settlements (Fig. 3). In Norway and Sweden, the majority of large towns with the highest concentration of permanent human settlements and high traffic-volume roads are located in the southern parts. Likewise, the farthest distance from the relic range and zones with lower annual wolverine reproduction goals are also in the south (Figs. 3 and S4). Thus, the combined effect of all these anthropogenic factors have probably limited the wolverine density distribution in the southern parts of the Scandinavian Peninsula. Nonetheless, the south represents the wolverine population’s expansion front and the observed latitudinal pattern may be also explained with the observation that wildlife population dynamics can differ considerably from the core areas (Swenson et al. 1998, Burton et al. 2010, Angert et al. 2020). With increasing human-made barriers to wolverine movement and dispersal (Aronsson and Persson 2018, Sawaya et al. 2019, Lansink et al. 2022), we expect the resulting population fragmentation will also play a major role in shaping the spatial distribution and dynamics of the Scandinavian wolverine population in the future.

As a measure of wild prey biomass availability, we included moose harvest density in our models (Table 1, Fig. S4). We estimated significantly higher wolverine densities in areas with higher moose harvest density, and this positive effect was more pronounced for males (Fig. 3). Wolverines are generally facultative scavengers and in many areas of Fennoscandia, they depend on slaughter remains from hunting and carcasses of prey killed by other top predators, including the Eurasian lynx *Lynx lynx,* wolf *Canis lupus,* and brown bear *Ursus arctos,* as well as animals dead from natural causes and roadkills (Van Dijk et al. 2008, Mattisson et al. 2011, Koskela et al. 2013, Aronsson et al. 2022). Moose occurs throughout the wolverine range in Scandinavia and moose carrion is an important food source for wolverines in many areas (Van Dijk et al. 2008, Mattisson et al. 2016, Aronsson et al. 2022), especially for breeding females (Koskela et al. 2013) and during winter (October - April) that overlaps with our study period. There is, however, considerable spatial and temporal variation in wolverine diet in Scandinavia, with reindeer as the most important prey for wolverines in some areas (Mattisson et al. 2016). Unfortunately, we were unable to find comprehensive and reliable data on the density of wild or semi-domesticated reindeer across the entire Scandinavian Peninsula to be considered for our study.

The positive effects of terrain ruggedness and the percentage of forest on wolverine density were significant for females only, while the average percentage of year-round snow appeared to only impact male density (Figs. 2 and S5). Traditionally, Scandinavian wolverines are not considered to be a forest-dwelling species, as they appear to select open and rugged terrain at higher elevations with snow, away from human activity (May et al. 2008, 2012, Rauset et al. 2013). Spring snow cover in particular is believed to be important for reproducing females as it determines denning suitability and offspring survival (Copeland et al. 2010, Mowat et al. 2020, Barrueto et al. 2022). However, in recent years, the Scandinavian wolverine population has expanded considerably into the boreal forest and has now colonized areas without persistent spring snow cover (Aronsson and Persson 2017). We chose the average year-round snow cover during the past decade not to specifically account for denning suitability for the wolverine, but as a measure of climatic niche suitability that may have shaped the wolverine’s density distribution today (Table 1). Terrain ruggedness and forest cover probably correlate with the degree of past persecution due to accessibility and history of land protection (Joppa and Pfaff 2009, Kerley et al. 2012) and the significance of these covariates for female wolverines may then reflect their affinity for high-quality habitat compared to males (May et al. 2008, 2012, Rauset et al. 2013, Aronsson and Persson 2018).

### Wolverines in the past, present, and future

Scandinavian wolverines have recovered from the brink of extinction and are now occupying a considerable portion of their historic range (Flagstad et al. 2004, Chapron et al. 2014, Gervasi et al. 2016, Aronsson and Persson 2017, Bischof et al. 2020). The effects of past impacts are nonetheless still clearly visible today, modulated, but not masked, by current environmental conditions and management regimes. The wolverine density in Scandinavia is shaped by human interests, while interacting with the history of local extinction. Wolverines are also impacted by other environmental covariates, several of which are directly or indirectly influenced by humans (e.g., prey base, climate conditions, and land-use). In an increasingly human-dominated landscape, the impact of humans on wolverines is likely to be even greater in the coming decades, further defining the state of the Scandinavian wolverine population. Despite the expansion of wolverines (Chapron et al. 2014, Aronsson and Persson 2017), an increasing human impact, if neglected, may therefore eventually again limit wolverines to the relic range that served as a refuge in the past.

## Data Availability Statement

Wolverine detections used in this study are available through the database Rovbase 3.0 at www.rovbase.no or www.rovbase.se. Data and R scripts of the spatial capture-recapture analysis will be deposited upon acceptance at: https://github.com/eMoqanaki.

## Authors’ contributions

R.B. and E.M. conceived and designed the study; H.B. provided the wolverine data and context on wolverine monitoring and management in Scandinavia; E.M. implemented the analysis with contributions from C.M., R.B., P.D., and H.B.; E.M. wrote the first draft and all co-authors discussed and contributed to the manuscript; all authors gave final approval for publication.

## Acknowledgments

We thank all contributors to the Scandinavian large carnivore monitoring database Rovbase 3.0. M. Tourani helped with the implementation of the variable selection approach and provided feedback on the figures. J. Kindberg provided access to the moose harvest data used in this study. S. Dey and S. Schowanek commented on an earlier version of the manuscript.

